# Two-step concentration of complex water samples for the detection of viruses

**DOI:** 10.1101/386060

**Authors:** Kata Farkas, James E. McDonald, Shelagh K. Malham, Davey L. Jones

## Abstract

The accurate detection and quantification of pathogenic viruses in water is essential to understand and reduce the risk of human infection. This paper describes a two-step method suitable for the concentration of viruses in water and wastewater samples. The method involves a tangential flow ultrafiltration step that reduces the sample volume of 1-10 l to approx. 50 ml, followed by secondary precipitation using polyethylene glycol 6000 that reduces the volume to 1-4 ml. For method validation, water samples were spiked with different concentrations of enteric viruses and viral recoveries in the concentrates exceeded 10% in all experiments. The method is suitable for water samples with high and low salinity and turbidity, allowing the accurate comparison of viral titers in a diverse range of water types. Furthermore, the method has the potential to concentrate other pathogens, e.g. bacteria or protozoa. Hence, the use of this method can improve the holistic assessment of risks associated with wastewater-contaminated environments.

## 1. Introduction

Enteric viruses (causing gastroenteritis) and other viral pathogens can be found in wastewater and in wastewater-contaminated surface and groundwater reservoirs. As the infective doses of these agents are low, concentration is needed to accurately quantify viruses in environmental waters and determine public health risks. A great variety of methods are available for water concentration for the recovery of viruses in wastewater and environmental water, however, many of those are not suitable and/or have not been validated for high volumes of water samples or for different water types. The most frequently used methods for the primary concentration of water samples are filtrations using electronegative (EN) or electropositive (EP) filters [1,2]. During EN and EP filtration, the water sample passes through the filter while the virus particles bind to the surface of the filter due to electrostatic forces. These methods have been shown to be suitable for the concentration of viruses in water, however, their use may be limited to low turbidity samples, due to filter clogging during filtration. Furthermore, the use of electronegative filters requires sample preconditioning (i.e. lowing sample pH), whereas electropositive filters may not be suitable for high salinity samples, and the elution of the virus particles from the filters may be difficult as well [1].

Tangential flow ultrafiltration (TFUF) has been used for the concentration of a wide range of water samples for the detection of various pathogens [1–5]. The main advantage of the TFUF approach is that during filtration the water flow takes place in parallel to the membrane, hence membrane clogging is less frequent compared to dead-end ultrafiltration and EN and EP filtration. In general, TFUF enables 40-200x concentration, and hence secondary concentration (filtration or precipitation) is often used to further reduce the sample volume [6–8].

This method describes an efficient, accurate and reproducible method for the concentration of enteric viruses in surface water (fresh and seawater), and wastewater (treated and untreated) samples (Figure 1). The recommended starting volumes are 10 l for surface water and 1 l for wastewater samples. The first step of the method is a TFUF step using a 100 kDa cut-off modified polyethersulfone membrane. As described by others, the efficiency of the elution of the viral particles from the membrane and the cleaning of the system was enhanced using sodium polyphosphate [4]. The final volume of the sample after TFUF is approx. 50 ml. In order to elute viral particles attached to solid matter in the primary concentrate, samples are further mixed with beef extract and then centrifuged. Then polyethylene glycol 6000 (PEG 6000) is added to the supernatant and viral particles are precipitated. These steps are based on a method for the elution and concentration of viral particles from sediment [9,10]. The resulting pellet contains the viral particles that can be eluted in phosphate saline buffer (PBS) and the solution can be stored at −80 °C. The concentrate may be subject to nucleic acid filtration, viral infectivity or integrity assays.

**Figure 1.**
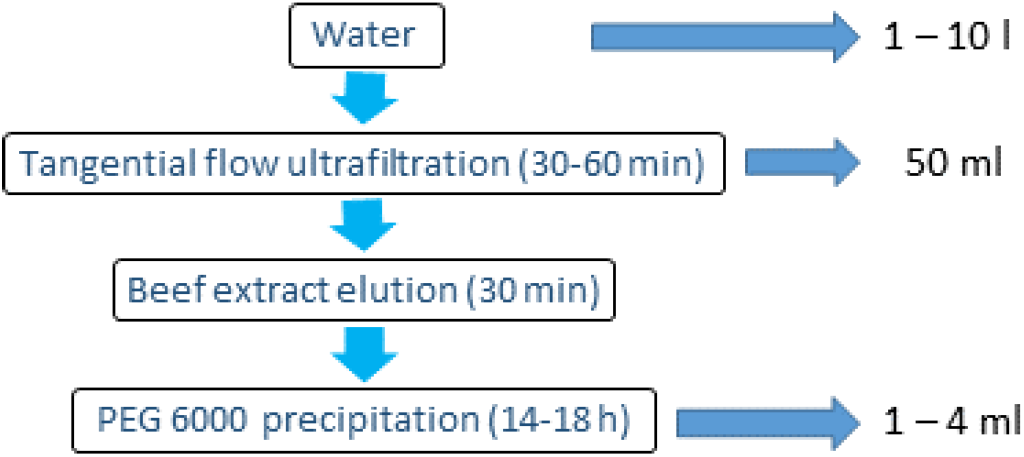
The stages of the tangential flow ultrafiltration-based two-step water concentration method for the detection and quantification of enteric viruses in water and wastewater.

## 2 Experimental Design

### 2.1 Materials

Sodium-polyphosphate (NaPP) / sodium hexametaphosphate (Sigma Aldrich, USA, Cat. No.: 305553)

Lab Lemco beef extract (Oxoid, UK, Cat. no.: LP0029)

Sodium nitrate (Sigma Aldrich, USA, Cat. no.: S8170)

Polyethylene glycol 6000 (PEG 6000) (Sigma Aldrich, USA, Cat. no.: 81255)

Sodium chloride (Sigma Aldrich, USA, Cat. no.: S7653)

Phosphate saline buffer (PBS), pH 7.4, (Gibco PBS tablets, Life Technologies, Cat. no.: 18912-014, Life Technologies, USA)

Virkon ^®^ solution (Lanxess, Germany)

20% ethanol (Fisher Chemical, #E/0650/17DF, Thermo Fisher Scientific, USA)

0.5 M HCl and 1 M NaOH for pH adjustment.

Optional: 30 µl mengovirus strain VMC0 (prepared according to ISO/TS150216-1:2013) solution with approx. 10^6^ mengovirus particles

### 2.2 Equipment

KrosFlo^®^ Research IIi Tangential Flow Filtration System (Spectrum Labs, USA, Cat no. SYR-U20-01N) or equivalent

100 kDa mPES MiniKros^®^ hollow fibre filter module (Spectrum Labs, USA, Cat. no.: S02-E100-05-N)

Silicone tubing #17 (Spectrum Labs, USA, Cat. no.: ACTU-E17-25N) or equivalent Centrifuge (2,500 xg and 10,000 xg at 4 °C)

Pocket sized pH meter (Ishiro, Japan, Cat no.: S2K992 or equivalent)

## 3. Procedure

### 3.1 Tangential flow ultrafiltration (Figure 2; Time of Completion: 2-4 hours)

#### 3.1.1 System wash

Wash system with 1 l 0.01% NaPP solution (0.1 g NaPP in 1 l deionized water) for 5 min (permeate closed) then leave the membrane in the solution for 30-60 min. Wash the membrane with the NaPP solution (permeate open) until the solution has been removed.

**Figure 2.**
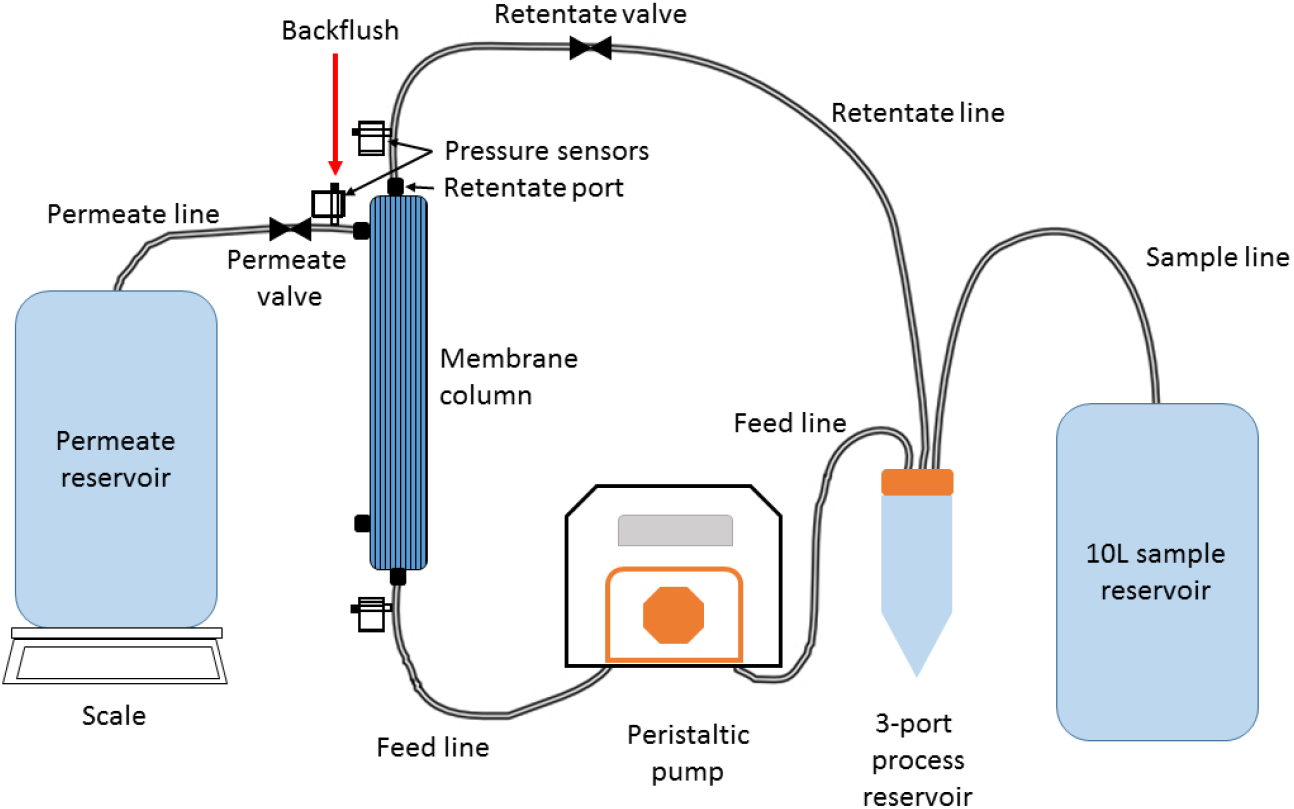
Schematic tangential flow ultrafiltration setup

#### 3.1.2 Sample filtration

**a) OPTIONAL STEP** Add approx. 10 µl mengovirus solution to the sample and mix. Save the rest of the mengovirus sample for control measurements.

b) Filter 10 l of surface water or 1 l wastewater at 1-1.6 l/min flow at 5 psi (0.3 bar; 30 kPa) pressure to achieve a permeate flow of 200 – 300 ml/min. Continue filtration until approx. 5 ml sample remains in the reservoir.

#### 3.1.3 Backwash, recovery

c) Set the flow to 680 ml/min with no pressure applied and circulate the concentrate for 5 min with the permeate clamp closed.

d) Stop pump, close penetrate and retentate valves.

e) Inject 20 ml 0.01% NaPP solution to penetrate pressure valve. Open retentate and wash with reverse flow.

f) Collect the concentrate from the system by introducing air through the retentate port. The final volume of the concentrate is approx. 50 ml.

#### 3.1.4 Membrane wash and storage

g) Wash membrane with 250 ml Virkon^®^ solution after each sample by circulating the Virkon^®^ solution in the system (permeate closed) at low flow (400-800 ml/min). In order to reuse membrane, immediately wash it with 150 ml 0.01% NaPP solution using the setup used for the Virkon wash. Repeat until solution in the process reservoir is clear. Leave the membrane in the solution for at least 10 min prior to reuse.

h) For long-term storage, wash the membrane with 50 ml 20% ethanol solution using the setup used for the Virkon wash. Repeat until solution is clear. Dissemble system and store membrane in 20% ethanol solution at 4 °C.

### 3.2 Secondary concentration (Time of Completion: 2.5 hours + overnight incubation)

#### 3.2.1 Virus elution

a)Add beef extract and NaNO_3_ to 50 ml concentrated water sample to reach final concentration of 3% w/v and 2 M, respectively. Adjust the pH to 5.5 using 0.5 M HCl.

b)Incubate at 50-90 rpm on ice for 30 min.

c)Centrifuge at 2,500 xg for 10 min, then transfer the supernatant to a new tube. Discard pellet. Adjust the pH of the solution to 7.5 using 1 M NaOH.

#### 3.2.2 Virus precipitation

d)Add PEG 6000 and NaCl to reach final concentration of 15% and 2% w/v, respectively. Mix to dissolve PEG 6000 and incubate at 4 °C for 14-18 h. 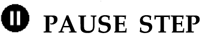 The solution may be stored at 4 °C for up to two days.

e)Centrifuge at 10,000 xg for 30 min at 4 °C. Discard supernatant. Dissolve pellet in 1-4 ml PBS (pH 7.4). The concentrate may be subject to infectivity/integrity assays or nucleic acid extraction flowed by real-time PCR quantification. Alternatively, viral nucleic acids can be extracted directly from the pellet.

**OPTIONAL STEP** For estimating method recovery percentile, RNA from 10 µl mengovirus solution should be extracted and quantified.

## 4. Expected Results

For pilot validation, 2 l of deionized water was spiked with of norovirus GII to reach the final concentrations of 10^6^, 10^5^, 10^4^, 10^3^, 10^2^ and 10^1^ genome copies (gc)/l in duplicates. Samples were concentrated using the two-step concentration method. Viral RNA was extracted from the pellet (final volume: 50 µl) and quantified using qRT-PCR (sample volume: 2 µl) as described in Farkas et al. [10]. High recoveries were observed in all norovirus concentrations (Table 1). The high deviations between replicates were result of the limitations of the qPCR method used for quantification. The limit of quantification (LOQ) was 200 gc/l and the limit of detection (LOD) was approx. 50 gc/l. The LOD and LOQ can be further lowered by the reduction of the RNA eluent volume and the increase of the sample volume in the qRT-PCR reaction.

**Table 1.** Norovirus recoveries and standard deviations (SD) in deionized water using the two-step concentration method.

| Norovirus concentration<br>(gc/l) | Recovery percentile<br>(SD) |
| --- | --- |
| $5.28 \times 10^6$ | 78 (43) |
| $4.72 \times 10^5$ | 121 (2) |
| $4.31 \times 10^4$ | 99 (11) |
| $3.03 \times 10^3$ | 91 (45) |
| $2.0 \times 10^2$ | 100 (0) |

For further validation, 10 l of surface water samples (river, estuarine and sea) in triplicates were spiked with known concentration of human enteric viruses (norovirus GII, sapovirus GI, hepatitis A virus and human adenovirus type 40) and with a mengovirus, which is often used as a process control for the extraction of enteric viruses from environmental matrices. The details of validation are described elsewhere [11]. Results showed that 10-100% recovery could be achieved in all sample types. In the same study, the usefulness of the method for wastewater samples was also investigated. Influent and effluent samples were taken in duplicates at four wastewater treatment plants. As the samples were expected to contain the target viruses, samples were processed without spiking. High viral concentrations were observed and the method showed great reproducibility as well. In subsequent samples spiked with mengovirus, at least 10% recovery was observed. No inhibition or cross-contamination between samples was observed. The method has also been successfully used for viral recovery from high volumes (50 l) surface water for metagenomics applications [12].

### Applications and recommendations

The method described above is suitable for the concentration of many different water samples and hence suitable for viral surveillance, contamination source tracking and to describe viral ecology. The TFUF method described here is suitable for the viral recovery from the mPES MiniKros^®^ hollow fibre filter (Spectrum Labs, USA), however, alternative recovery buffers may be used with different membranes. Furthermore, as the membrane used for the TFUF has a 100 kDa pore size, the method can potentially co-concentrate other microbes and protozoa as well, and hence the TFUF step of the concentration method enables the accurate description of microbial quality of a sample. In the current study, viral nucleic acids were directly extracted from the concentrated water samples, however, the concentrates are suitable for viral infectivity and capsid integrity assays as well [11]. For direct nucleic acid extraction, the resuspension of the PEG precipitate is not necessary; nucleic acids can be extracted from the pellet. When PCR-based approaches are used for the quantification of viruses in the concentrate, the use of robust extraction and amplification methods is recommended, as organic matter that may interfere with the enzymes used for amplification are co-concentrated with viral particles. The addition of process control (e.g. mengovirus) to each sample to determine method efficiency is highly recommended.

## Author Contributions

Conceptualization and Methodology, All; Validation, Investigation, K,F.; Writing-Original Draft Preparation, K.F.; Writing-Review & Editing, All; Funding Acquisition, D.J.

## Funding

This work was funded by the Natural Environment Research Council (NERC) and the Food Standards Agency (FSA) under the Environmental Microbiology and Human Health (EMHH) Programme (NE/M010996/1).

## Acknowledgements

The authors would like to thank Lindsay Smart (Spectrum Labs, UK) for his assistance in the method development for water concentration.

## Conflicts of Interest

The authors declare no conflict of interest.

